# Harmonizing Peak Matching Between Multidimensional NMR Spectra

**DOI:** 10.64898/2026.02.21.707192

**Authors:** Anthony C. Bishop, Kyle Mimun, Weimin Tan, Taylor R. Cole, A. Joshua Wand

**Affiliations:** Department of Biochemistry & Biophysics, Texas A&M University, College Station, Texas 77843 USA

**Keywords:** heteronuclear NMR, cross-peak matching, Bayesian statistics, automation, confidence estimation, peak tracking

## Abstract

A common task in analysis of multidimensional NMR is the mapping of cross-peaks from one spectrum to another. In some situations, only a subset of coordinates of cross-peaks are to be matched such as in comparing corresponding dimensions of triple resonance assignment spectra. In other cases, the sample is perturbed in some way, and the task is to map corresponding cross-peaks that potentially change in any and all dimensions. This exercise is commonly done “by hand” and is both time-consuming and subjective. It is difficult for an individual to consider all possible interpretations of deviations between two multidimensional NMR spectra. Furthermore, in regions of high degeneracy, cross-peak matchings between spectra can be ambiguous. Ideally, a mapping algorithm should reflect a confidence of its matchings to indicate the reliability of proposed matchings and should do so while harmonizing all potential mappings in the spectrum. Critically, this process should be automated using criteria that are both well-defined, consistent and provide a measure of reliability. Here we develop a novel model describing the distribution of apparent deviations between cross-peaks of two multidimensional NMR spectra to random noise and true deviations to harmonize the cross-peak matching. Bayesian inference is employed to provide confidence estimates. The resulting algorithm (pHarmony) is tested in a variety of ways such as mapping between pairs of triple resonance spectra and between two- and three-dimensional heteronuclear NMR spectra generated by a fragment-based ligand discovery screen.

## Introduction

Analysis of multidimensional NMR spectra invariably requires the mapping of individual cross-peaks arising from a set of common spin resonances. In some cases, the mapping is a subset of nuclei such as the comparison of amide N-H correlations in, for example, HNCO and HN(CA)CO spectra. In other cases, the entire spin system being correlated is mapped to a corresponding cross-peak arising in perturbed spectrum. Chemical shift perturbations can arise from changes in physical environment (e.g., temperature, pressure, ionic strength, etc.), variations of between protein samples that are ostensibly the same or the addition of a ligand such as might occur in a ligand discovery campaign. Tracking the movement of cross-peaks provides valuable biophysical information on the system of interest. Often cross-peaks are identified (assigned) in a reference spectrum and “matched” to their corresponding cross-peaks in other spectra, most often as a part of a manual workflow. Should the number of spectra and number of involved cross-peaks be small, manual peak picking and matching is not a significant bottleneck. More commonly, the number and complexity of spectra to be compared, such as those encountered in a triple-resonance assignment project or an NMR based fragment-based drug discovery screening campaign, becomes unwieldy and automation of the process is desirable. Furthermore, tracking cross-peaks between spectra can be ambiguous and there is great utility in having an estimate of confidence in a given cross-peak match. Though numerous automated peak picking algorithms have been proposed ^1-3^ relatively few algorithms exist for the purpose of cross-peak matching and generally do not provide estimates of confidence for the proposed matches nor operate on spectra of arbitrary dimensionality. Current cross-peak matching algorithms are also designed to work on manually picked peak lists, where the peak picking was done with expert reliability and consistency and not with automated algorithms which may fail to consistently pick cross-peaks across spectra in regions of high degeneracy. To address these limitations, we have developed the cross-peak matching algorithm pHarmony, which can be used on spectra of arbitrary dimensionality, provides a rigorous confidence score for its proposed matches, and is robust when handling automatically picked peak lists.

### Theory

Here we outline the pHarmony algorithm which is designed to harmonize the mapping of cross-peaks of one spectrum (the reference) with the cross-peaks of another spectrum (the target). A graphical summary of the probability model employed is provided in Fig 1. The various elements of the model are described below.

**Fig. 1.**
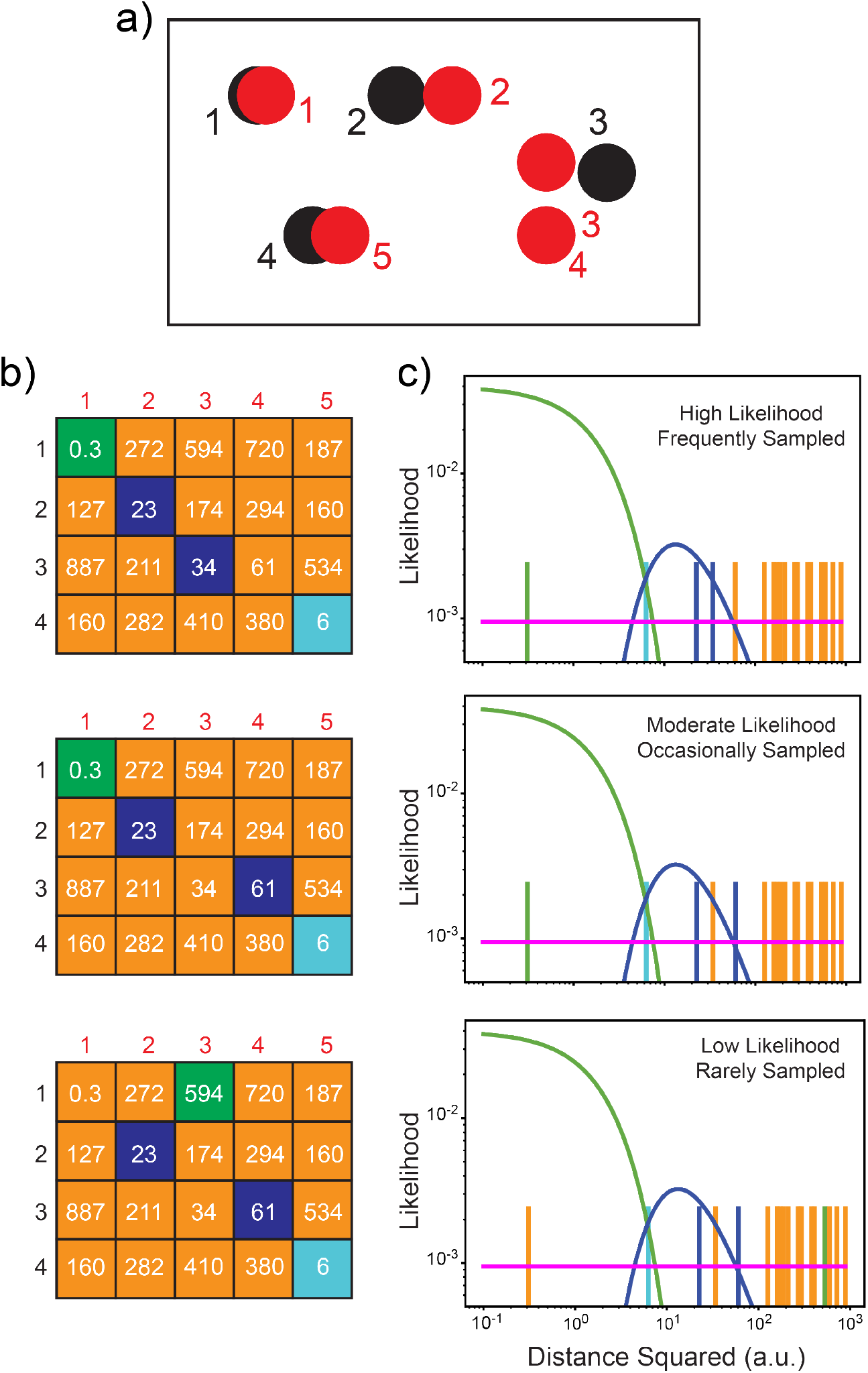
Probability model underlying the pHarmony algorithm. Panel a) Schematic of an overlay of a reference spectrum (black) and target spectrum (red). Numbers correspond to indices in Panel b. Panel b) Examples of sampled matching matrices with the value of the squared normalized distance between the corresponding peaks (arbitrary units) in each cell. Orange tiles correspond to cross-peak pairs that were not matched. Green, blue and cyan tiles correspond to cross-peaks that were matched with distances signifying random noise, arising from chemical shift perturbation (CSP) or with similar probability of arising from random noise or CSP, respectively. High likelihood (top), moderate likelihood (middle) and low likelihood (bottom) matching matrices are shown as examples. Panel c) Likelihood plots corresponding to b) as a function of squared normalized distance for each matching matrix at a fixed distribution parameterization in log-log space. Green lines correspond to a ×^2^ distribution with two degrees of freedom to model squared normalized distance arising from random noise between matching cross-peaks. Blue lines correspond to a Fréchet distribution of fixed parameterization modeling squared normalized distance arising from true CSPs between matching cross-peaks. Horizontal magenta lines correspond to the approximately uniform distribution in two-dimensional space of distances between cross-peaks that do not match. Each vertical bar corresponds to a cell in the matching matrix and its color indicates the classification made (see Panel b). Distances are more likely to be classified into a given distribution according to the density of that distribution at a specific squared normalized distance. In principle, the frequency at which each matching matrix will appear is proportional to its likelihood (see sampling the posterior distribution).

The problem of matching *R* reference spectrum cross-peaks to *T* target spectrum cross-peaks can be described as finding the *R* x *T* matrix *M*. Entries in *M* are either a one (1) or a zero (0) where a value of 1 at position *r, t* describes a match between the reference peak *r* and the target peak *t*. A zero indicates not a match. We constrain the problem domain to circumstances where cross-peak movement is due to either spectral noise or fast-exchange phenomena such as that which generally occurs with weak ligand binding. Thus, *M* is subject to the constraint that there can be at most a single “1” in any given row and column. This enforces exclusivity (i.e., a reference cross-peak can only match at most one target peak and vice versa). We permit *M* to encode circumstances where cross-peaks have no match (i.e., rows and columns that are filled with zeros) to account for the possibility that a cross-peak present in one spectrum has no match in the other. This provides tolerance for the situation where a peak may have disappeared for physical reasons (e.g., due to intermediate exchange broadening) as well as making it robust to errors in upstream data analysis (e.g., erroneous cross-peak picking).

Effectively, in current manual approaches, the finding of *M* leads directly to subsequent analysis. However, it is often difficult to be certain that any proposed matching matrix is equivalent to the true matching matrix *M*. For example, it is often the case that a spectral perturbation (e.g., due to ligand binding) could induce chemical shift perturbations (CSPs) in multiple cross-peaks in a crowded region. Without additional information, assignment of matches is ambiguous even for the experienced spectroscopist. Thus, instead of only proposing a single matching matrix as the most likely *M*, to describe the matching, it is more valuable to model the matching matrix *M* itself as a random variable and determine its Bayesian posterior probability distribution.

If a model is defined for calculating the likelihood of any possible *M* (defined as *M*_*i*_), it becomes possible to apply Bayes’ rule to calculate its posterior probability:

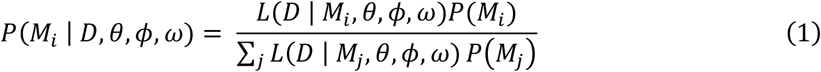

*L* is a function that describes the likelihood of observed peak perturbations given a possible matching matrix *M*_*i*_ and parameters *θ, ϕ, ω. P*(*M*_*i*_) is the prior probability of any possible *M*_*i*_. We assume that the prior distribution of *M*_*i*_ is uniform and can thus ignore its contribution to posterior probability essentially assuming that *P*(*M*_*i*_) is equal for all *M*_*i*_.

Explicitly calculating the partition function (i.e., the denominator of equation 1) would require calculating the likelihood of all possible *M*_*i*_. Given that the number of possible *M*_*i*_ scales combinatorially with the number of cross-peaks, an explicit calculation of the partition function is intractable for even a dozen or so cross-peaks. Furthermore, the values of the likelihood function parameters (i.e., *θ, ϕ, ω*) are not known *a priori* and need to be determined.

### Feature extraction

Perhaps the most intuitive spectral feature to use to match cross-peaks between reference and target spectra is the chemical shift distance between them. The pHarmony algorithm calculates an *R* x *T* normalized distance squared matrix *D* between all possible pairs of reference and target cross-peaks. Calculation of an uncertainty-normalized distance squared between reference peak *r* and target peak *t* utilizes the Equation 2:

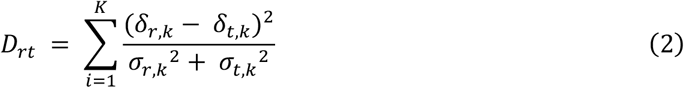

Where *δ*_*r,k*_ and *δ*_*t,k*_ are the chemical shifts of cross-peaks *r* and *t* respectively, along spectral dimension *k* and *σ*_*r,k*_ and *σ*_*t,k*_ are the estimated uncertainty in the position of cross-peaks *r* and *t* respectively along dimension spectral dimension *k*. A separate uncertainty for each dimension in each spectrum is used for their respective peak uncertainties. Elements of D are referred to below as normalized distances squared.

### Classification and Likelihood Model

The algorithm models the problem of matching *r* reference spectrum cross-peaks to *t* target spectrum cross-peaks as an unsupervised classification problem of the normalized pairwise squared distances. The matching matrix *M* can be thought of as a classification of each element in *D*. Each element of a possible matching matrix *M*_*i*_ classifies its corresponding element within *D* to one of two possible categories: (1) the two cross-peaks defining the distance do not match or (2) the two cross-peaks defining the distance do match, subject to the previously mentioned constraints of at most one-to-one matching of reference to target cross-peaks. Of the distances that correspond to a match, we then classify these further into matching cross-peaks with a distance that arises due to random noise or due to an actual CSP (i.e., a real difference). This second layer of classification is described by matrix *C* with dimensions *R*× *T*. Each element *C*_*rt*_ is either a zero or a one depending on whether the corresponding entry in *D* is classified as arising due to only random noise or due to a CSP, given the corresponding cross-peaks match. Unlike *M*_*i*_ where the probability that each element is a one or a zero is highly dependent on the other elements, it is assumed that the probability that the value of any element *C*_*rt*_ is independent of all other elements in *C*. Given a pair of matched cross-peaks, the probability that the matched pair arose from a CSP or noise is independent of whether any other pair of matched cross-peaks arose from a CSP or noise. Each of these categories in which a distance *D*_*rt*_ can be placed (non-matching pair, matching pair perturbed by noise, matching pair with a CSP), will have a defined likelihood function that is used to calculate the likelihood of an observed *D*_*rt*_ given that the corresponding peak pair is assigned to that category. The likelihood functions of each category are defined below:

### Matching cross-peaks perturbed by random noise (*M*_*rt*_ = 1, *C*_*rt*_ = 0)

For the elements *D*_*rt*_ that arise between matching cross-peaks differing in position from random noise, the positions of the corresponding cross-peaks *r* and *t* are modelled as multivariate Gaussian random variables centered at the observed chemical shifts (e.g., *δ*_*r,k*_ and *δ*_*t,k*_) with standard deviations given by the uncertainties (*σ*_*r,k*_ and *σ*_*t,k*_) for each spectral dimension *k*. Given this definition, *D*_*rt*_ can be modelled as a *X* ^2^ random variable of *K* degrees of freedom where *K* is the number of spectral dimensions. We thus model the likelihood function for the matching perturbed by noise case as the *X* ^2^ PDF:

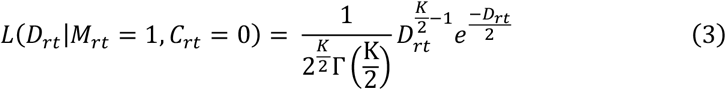

### Matching cross-peaks perturbed by a CSP (*M*_*rt*_ = 1, *C*_*rt*_ = 1)

Elements *D*_*rt*_ that correspond to matching cross-peaks and arose from CSPs are modelled as IID Fréchet random variables. Unlike the *X*^2^ random variable that has a parameterization known *a priori*, the optimal parameterization for this distribution needs to be learned. The likelihood function for this case is given below

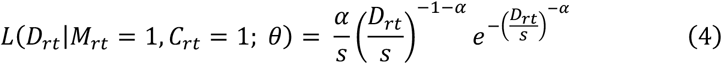

Where *α* and *s* are the shape and scale parameters for the distribution (collectively referred to as the parameter vector *θ*).

### Distances between non-matching cross-peaks (*M*_*rt*_ = 0)

In addition to the two distributions that model elements *D*_*rt*_ between matching cross-peaks, we introduce a third function that defines the likelihood for the elements *D*_*rt*_ between cross-peaks that *do not* match. The form of this likelihood is taken as the following:

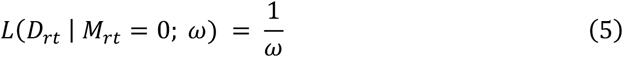

This is a pseudo-likelihood based on the observation that the distribution of normalized square distances of between non-matching peaks is approximately uniform at moderate distances from the peak (see Figure 3 below) i.e., the function is constant with respect to *D*_*rt*_. The parameter *ω* is calculated from a user-supplied input based on the furthest distance the user would expect a peak to move before considering it equally likely that the original peak disappeared and a different peak appeared. It functions as a threshold that guarantees the likelihood of a pair of peaks not matching will exceed the likelihood of the peaks matching at a specified distance. (See the discussion below on expectation-maximization for more detail)

Using the three above components we can express the total likelihood as a hierarchical mixture model of the three categories:

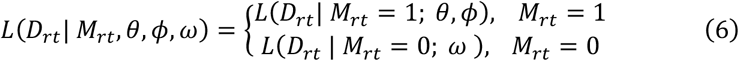

Where *L*(*D*_*rt*_| *M*_*rt*_ = 1; *θ, ϕ*), the matching likelihood, is the mixture of the *X*^2^ and Fréchet distributions, which describes the pdf of *D*_*rt*_ given that the corresponding cross-peaks match. It is defined as:

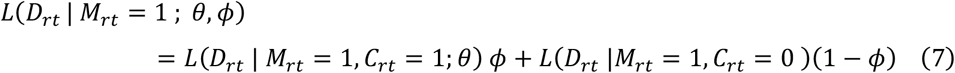

Where *ϕ*, bounded on [0,1], describes the relative contribution of the two components to the matching likelihood. Together this provides a model allowing the evaluation of the likelihood *L*(*D*_*rt*_ | *M*_*i,rt*_, *θ, ϕ, ω*) for any given set of *M*_*i*_, *θ, ϕ, ω*. Assuming that the likelihood *L*(*D*_*rt*_ | *M*_*rt*_, *θ, ϕ, ω*) is independent among all *D*_*rt*_ then the likelihood of the entire matrix *D, L*(*D* | *M*_*i*_, *θ, ϕ, ω*) can be calculated as the product of all *L*(*D*_*rt*_ | *M*_*i,rt*_, *θ, ϕ, ω*) for every combination of r and t.

### Sampling the Posterior Distribution *P*(*M*| ***D, θ, ϕ, ω***)

We have now defined a model for describing the likelihood of any given matching matrix. Given a set of parameters and the matrix *D*, we want to determine the posterior distribution of *M*. As noted previously, an exact evaluation of the distribution’s pdf requires computing the likelihood of every possible matching matrix. To avoid this computationally prohibitive step, the matching matrix distribution is sampled using a sequential Monte Carlo sampler ^4^, designed such that matching matrices are sampled with the probability described by Equation 1. The sequential Monte Carlo (SMC) sampler treats sampling a matching matrix as a series of decisions. First, a decision likelihood matrix (*S*) of dimensions *R x*(*T* + 1) is constructed, where the likelihood at position *r, t* corresponds to the decision of matching reference peak *r* to target peak *t* and where position *r,T+1*, corresponds to the likelihood of reference peak *r* having no match in the dataset. The likelihood of the decision at position *r,t* (*S*_*rt*_)is calculated by taking the product of the matching likelihood of peak *r* with target peak *t* with the product of all of the non-matching likelihoods with every other target peak. For the case of matching reference peak *r* with no target peak, the decision likelihood (*S*_*r,T*+1_) is the product of the non-matching likelihoods of the reference peak with all target cross-peaks i.e.,

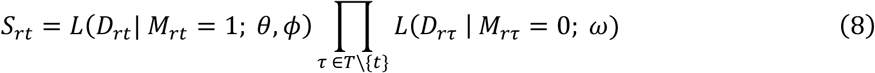

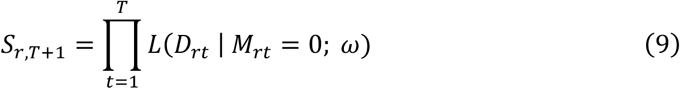

The rationale behind calculating the decision likelihood in this manner is if reference peak *r* is matched to target peak *t*, then by the previously stated constraints *r* must also *not* match every other target peak. Similarly, the likelihood of not matching any target peak is the product of the non-matching likelihoods for each target peak.

Additionally, the constraints of our problem mean that the decision likelihood matrix itself will change as future decisions become unavailable. Once a decision is made for reference peak *r*, the likelihoods for all decisions in row *r* (*S*_*r*,∗_) are set to 0. In other words, once a decision for a row has been made, another decision cannot be made for that row. Likewise, if the decision *S*_*rt*_ is made, then the likelihood matrix for the entire column *t* (*S*_∗,*t*_) will be set to zero as a target peak cannot be matched twice. The exception is the column (*S*_∗,*T*+1_) which will always have a non-zero likelihood for all reference cross-peaks that have not had a decision yet made (i.e., any number of reference cross-peaks can be labelled as not matching any target peak).

### The SMC target and proposal distributions

Initially the SMC matching algorithm was implemented as described^4^. However, it was determined that the simplified target distribution was insufficient to adequately match cross-peaks and, following Jun et al^4^, employed a proposal distribution that is calculated to match the target distribution (Equation 1) as closely as possible. Since each entry in the decision likelihood matrix describes the likelihood of an entire row of the matrix *D*_*rt*_ (i.e., each decision in row (*S*_*r*,∗_) represents a different possible *M*_*r*,∗_). The likelihood function *L*(*D* | *M*_*i*_; *θ, ϕ, ω*) of the target distribution can be reparametrized as the product of a set of decisions:

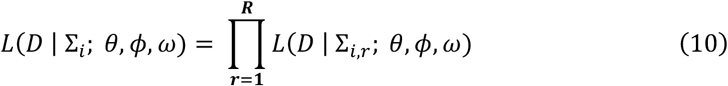

Where ∑_*i*_ refers to any complete, valid decision set (i.e., a single decision made for each reference peak) and ∑_*i,r*_ represents a valid decision for reference peak *r*. To emphasize, the likelihood function presented here is the same as defined earlier; the difference is the matching matrix is now represented by a sequence of decisions. This implies that any given ∑_*i*_ maps exactly one to one with a possible matching matrix *M*_*i*_. We can then completely reparametrize equation 1 as:

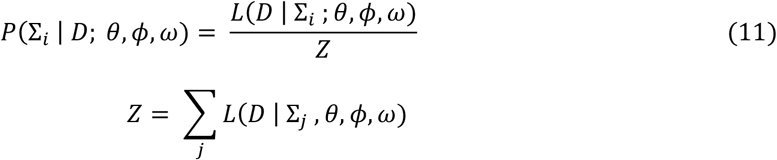

The summation over j involves summing the likelihoods of all possible complete decision sets. Additionally, the prior probability terms have been dropped as we assume that all prior probabilities of all valid decision sets are equivalent to one another.

In SMC sampling, when drawing a sample of decision sets of size *N*, all members of the sample are drawn simultaneously. The probability of sample member *i* sampling decision set ∑_*i*_ is defined by a proposal probability *Q*(∑_*i*_ | *D*; *θ, ϕ, ω*). Because the proposal distribution does not necessarily equal the target distribution, each member *i* in the sample is given an associated weight which is defined as the ratio of the target and proposal probabilities.

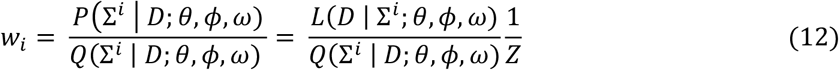

A new sample is drawn by randomly drawing members from the current sample (with replacement) with a probability proportional to their weight. The probability of drawing member *i* from the sample is equal to:

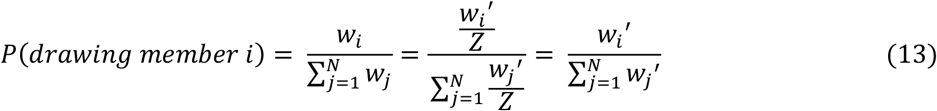

As the partition function *Z* has no bearing on the resampling probability, *Z* can be dropped from the calculation of the sample weights. Thus, weights can be calculated as the ratio of the likelihood of the target distribution to the proposal distribution probability.

Resampling guarantees that the drawn sample follows the target distribution. However, if the proposal distribution deviates significantly from the target, the weight can become concentrated in just a handful of sample members leading to high sample variance. Because of this, it is helpful to resample before all decisions have been made. This allows sample members that have “fallen off track” and are exploring low probability regions of the target distribution to be discarded. Resampling after a subset of decisions is possible given the ability to factorize our target distribution:

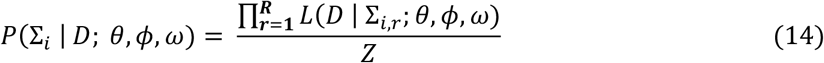

Which can be rewritten as

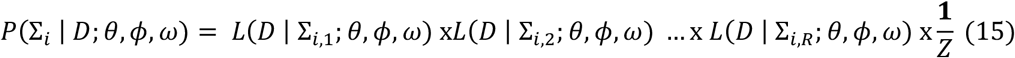

We can also define a target distribution over a set of partial matches

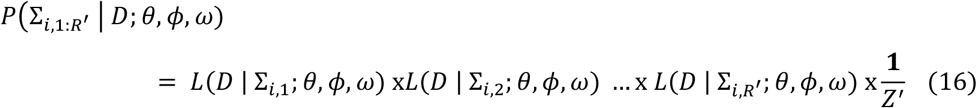

where 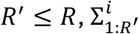 is the partial set of decisions over the first *R*^′^ reference cross-peaks and *Z*^′^ is the partition function over the decision set for the first *R*^′^ reference cross-peaks. The proposal distribution can similarly be decomposed in the following manner.

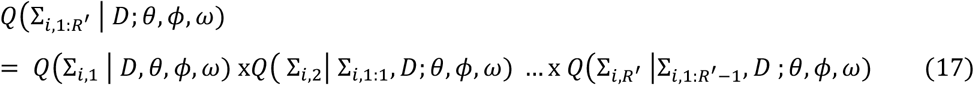

Each factor in the proposal distribution is essentially the proposal probability of the decision for each reference peak. It is important to note that the proposal probability has a conditional dependence that the target probability does not – i.e., the proposal probability distribution over the possible decisions of row *r* is dependent on all preceding decisions. This accounts for the fact that a target peak that was matched during a previous decision is no longer available for matching. Thus, the weight for sample member *i* after sampling decisions for the first *R*^′^ rows is:

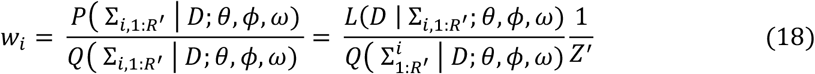

And the probability of sampling member *i* during resampling is

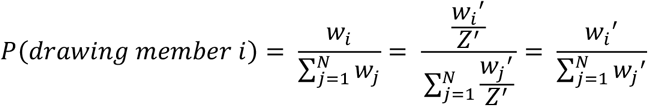

As before, the partition function *Z*^′^ can be ignored thus permitting the weights to be calculated as the ratio of the target likelihood of the partial decision set to the proposal probability of the partial decision set. This enables accurate resampling at any step.

### Computing the proposal probability

One possible proposal probability for step *r* is

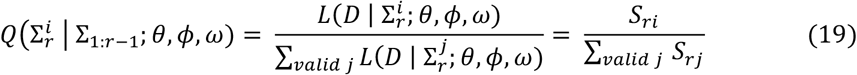

Where the superscripts *i* and *j* index a possible decision available to reference peak *R*^′^ (i.e., that haven’t been removed from consideration by earlier decisions). This proposal distribution however often leads to extremely uneven weights across the members of the sample and very high sample variance. The issue lies in the tendency to create seemingly good matches in the earlier decisions which preclude better matches that could have been made to later reference cross-peaks. Thus, a good proposal distribution needs to account for the effect of its current decision on the likelihood of all possible future decisions. The ideal proposal would be as follows:

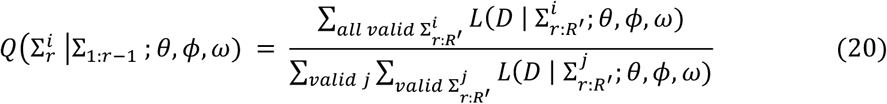

Where 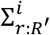 refers to a partial set of decisions which matches reference peak *r*^′^ with the target peak at index *i* (or the non-matching decision at *i = T+1*) and any possible decision for each of the subsequent rows up to row *R*^′^. Calculation of the exact proposal is intractable as the denominator of this equation, for the first decision (*r* = 1, *R*^′^ = *R*), is the full partition function shown in equation 1. However, it is often the case that the majority of the sum over the possible likelihoods of partial decision sets 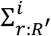 is contained in relatively few of the possible partial decision sets. Furthermore, a decision made at reference peak *r* will not substantially influence the probability of decisions made at all other reference cross-peaks, just a subset of them. Thus, if the reference cross-peaks that substantially affect each other’s proposal distributions were ordered such that they were close in the decision sequence, then the bulk of the *partial* partition function would be accurately estimated by calculating considering only decisions at the “most relevant” subset of subsequent rows. In effect, the goal is to group the reference peaks in the decision sequence such that highly correlated reference peaks are close in sequence and to identify groups of reference peaks where decisions made for reference peaks in one group do not substantially affect the probabilities of decisions made for reference peaks in another group.

### Reference Peak Ordering

Reference cross-peaks were ordered by first clustering the reference cross-peaks together via hierarchical clustering and then choosing an ordering that keeps the nearest neighboring cross-peaks close in sequence. A distance matrix for the purpose of clustering reference cross-peaks is calculated in the following way. First, for each reference peak, a subset of the target cross-peaks is chosen as matching candidates for that target peak. Any target cross-peak with a matching likelihood that was less than the no match likelihood was eliminated from its candidate set. The no match decision was kept as a candidate for each reference cross-peak. The likelihoods of the candidate decisions were then normalized to a probability. A target peak was kept as a candidate if its probability was within a factor of 20 of the largest probability target peak for that reference cross-peak. This was done to eliminate target cross-peaks from a reference cross-peak’s candidate set in cases where the target had a much higher probability of matching a different reference cross-peak. The number of overlapping candidate target cross-peaks between two reference cross-peaks was computed for all reference cross-peak pairs. For a common target cross-peak to be counted as overlap between two reference cross-peaks, the target must be present in both sets, and the ratio of the probability for the two reference-target cross-peak matches must not differ by more than a factor of 20. This ensures that overlap is only counted if there is a reasonable chance that the same target peak is matched to either reference peak. In other words, two reference cross-peaks can only be considered as “competing” for a target peak if the probability doesn’t seem to overly favor one reference peak over the other.

After determining the number of overlapping candidates between each pair of reference cross-peaks, a distance between each pair of reference cross-peaks was calculated by taking the multiplicative inverse of the number of overlapping target cross-peaks.

Reference peak pairs with no overlap were assigned a distance of infinity. From this distance matrix, a weighted shortest path matrix was computed by finding the shortest path between all pairs of reference cross-peaks using the Floyd-Warshall algorithm ^5^. For example, if the shortest path from reference peak A to C was via a third reference peak B, the path distance between A and C was set to the sum of the distance between A and B and the distance between B and C. The shortest path matrix was then used by the scikit-learn^6^ AgglomerativeClustering class, using the single-linkage clustering algorithm and a distance threshold of 1.0. This guarantees any two reference cross-peaks that have a path of non-infinite distance between them will belong to the same cluster. Using the clustering results, a linkage matrix defining a dendrogram was calculated which was passed to the scipy^7^ optimal leaf ordering algorithm, the results of which became the decision order for the reference cross-peaks. This resulted a reference peak order with independent clusters of reference peaks, where highly correlated reference peaks were placed proximally in the decision sequence within a cluster.

### Beam Search

To compute the proposal distribution a beam search strategy ^8^ was used to estimate the partition function of over the remaining decisions within a cluster of reference peaks. For reference cross-peak *r* a beam search of width 1000 is performed over all of the remaining reference cross-peaks in that cluster. From a given candidate decision within *r*, a beam is sent to every candidate at reference *r+1*. The beams then split such that each *r+1* candidate sends a beam to each candidate decision at *r*+2. With the repeated branching the number of beams grow exponentially. Should the number of beams exceed 1000 at any time only the 1000 beams with the highest likelihoods are retained, with the likelihood of each beam being the product of the likelihoods of the decisions it visits. This continues until the last reference peak in the cluster is reached. The likelihoods of all of the beams are then summed together, providing the proposal weights for that decision at reference peak *r*. This is repeated for each possible decision in *r*. The weights for each decision are normalized over the decisions to yield the proposal probability distribution for the decisions of reference peak *r*.

### Resampling and sample size determination

Once a decision for the last reference peak in a cluster has been made, the effective sample size (ESS) ratio (defined as the ratio of the effective sample size to the sample size *N*) is calculated.

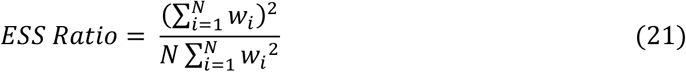

The ESS ratio is indicative of how well the proposal distribution matched the target distribution across the sampled decisions with a value of 1.0 indicative of perfect agreement and 0 being very poor agreement. If an ESS ratio of 0.2 or larger is measured the sample is resampled by stratified resampling^9^ where a new sample is drawn from the current sample (with replacement) such that the probability of drawing a member of the sample is proportional to its sample weight. If the ESS ratio is less than 0.2, indicative of poor agreement between proposal and target distributions, all decisions made on the reference peaks in the current cluster are discarded and the algorithm returns to the first cross peak in the cluster. The maximum width of the beam search is increased by a factor of five and the algorithm proceeds as normal. If the maximum beam search width becomes too large such that an internal memory threshold is exceeded, the program errors out. This occurrence seems quite rare in practice and has only been seen in cases of highly unrealistic peak lists (e.g. grossly incorrect peak picking).

At the beginning of the algorithm, sample size *N* is initialized to the number of reference cross-peaks. After sampling the posterior distribution, the frequency of each reference-target cross-peak match is calculated, and a second sample is drawn. The frequencies of each reference-target match are compared. If the difference in frequency for any reference-target pair is greater than 0.1, *N* is increased by a factor of two and the sampling is repeated.

### Determining *θ, ϕ*, and *ω* via Expectation-Maximization

Though we have developed a method for sampling the posterior distribution of *M*, this distribution is parameterized by *θ*, ϕ, and *ω*, the true values of which are not known *a priori*. It is possible to treat these parameters in a fully Bayesian fashion and determine their posterior distributions; however, as these quantities are of indirect interest, it should suffice to determine the maximum likelihood estimates (MLE) and the maximum *a posteriori* (MAP) estimate for *θ*, ω and the mixture model weight ϕ, respectively. In expectation-maximization optimization, the posterior distribution of matching matrices is sampled and the sampled matching matrices are then used as inputs to the target likelihood function to optimize the parameters. The algorithm begins with initial estimates for *θ*, ϕ, and *ω*. The initial estimate of *θ* is chosen by collecting the smallest value *D*_*rt*_ at each value of *r*, eliminating all values < 3 and finding the MLE for the Fréchet distribution parameters with respect to these elements, bounded such that the shape parameter is greater than or equal to 1. For the mixture weight parameter ϕ which will ultimately be estimated via MAP, we define a prior distribution using a Beta distribution and use its mean as the initial estimate for the MAP value. The parameters of the Beta distribution (*α, β*) are influenced by the expected characteristics of the matching problem. The user provides a rough estimate of what fraction of reference cross-peaks are expected to be perturbed by CSPs. The parameters of the beta distribution are calculated such that its mean will be equal to that value. The standard deviation of the distribution is set (by default) to two times that of the distribution mean but can be scaled with a user-supplied variance scaling factor. If either *α* or *β* are calculated such that it is less than 1, both parameters are rescaled such that the mean of the beta distribution is maintained, but both *α* and *β* are greater than or equal to 1 which is necessary for a proper MAP update during EM. For ω, the user supplies an estimate for the largest CSP expected. Using *θ* and ϕ, the matching likelihood at the normalized distance squared corresponding to the largest CSP is calculated. ω is calculated such that the matching and the non-matching likelihoods are equal at the largest expected CSP.

### Expectation step

With initial estimates for *θ*, *ϕ*, and *ω*, we can then use the sampling strategy described previously to generate a set of matching matrices, ℳ. The posterior probability of matching between any two cross-peaks n and m can be estimated using ℳ by calculating the frequency at which a match between each reference and target peak occurs in the sample:

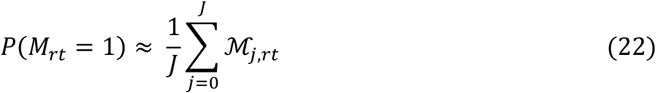

Where ℳ _*j,rt*_ refers to the element at row *r* and column *t* for, the *j*th matching matrix in sample ℳ. In addition, we can calculate the posterior probability describing that any *D*_*rt*_ arose from a CSP given that that reference peak *r* and target peak *t* match. This is calculated in the following manner:

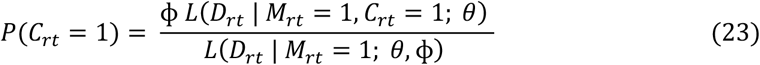

### Maximization step

With the posterior sample of matching matrices and the posterior CSP probabilities, MAP values for *ϕ*, and *ω* can be updated. In effect, *ϕ* is the parameter of a categorical distribution (with a range over two categories). Since the Beta distribution prior is conjugate with respect to the posterior distribution of *ϕ*, it can thus be used within a closed form expression to provide a regularized update:

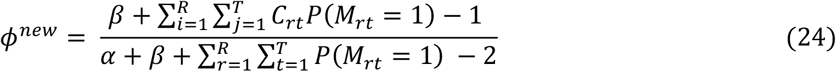

Where *α, β* are the parameters of the prior distribution of *ϕ*.

In the case of *θ* there is no closed form expression allowing for a direct calculation of an updated MLE. Instead, we approximate the solution numerically by finding *θ* with the maximum likelihood under the posterior distribution of matching matrices. This can be done by optimizing theta such that the product of the sampled matching matrix likelihoods is maximized.

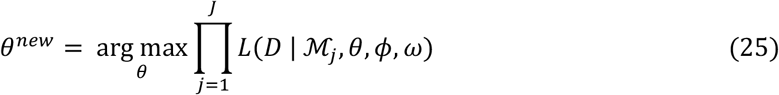

with the goal of finding *θ* with the maximum likelihood with respect to the sample. With *θ* and *ϕ* updated, *ω* is updated by calculating the value of *ω* that makes the matching likelihood and non-matching likelihoods equivalent at the maximum expected CSP.

With these updated parameters (*θ, ϕ*, and *ω*), a new round of sampling the posterior distributions can commence, followed by an additional round of parameter updates, and continuing until parameter convergence. Once the converged parameters have been identified, the posterior distribution of matching matrices can be sampled and posterior probabilities of individual matches calculated.

## Materials & Methods

### NMR Spectroscopy

Uniformly ^15^N-^13^C-labeled human IL-1β was expressed recombinantly in *E. coli* BL21(DE3) cells and purified essentially as described elsewhere^10^. IL-1β was encapsulated in LDAO/10MAG reverse micelles^11^ in hexane with 2% D_14_ – hexane at an aqueous concentration of 5 mM as described previously ^10^. NMR data were acquired at 25°C on an 800 MHz four channel Bruker NEO NMR spectrometer equipped with a cryogenically cooled probe. Three-dimensional HNCO experiments were collected with 512 × 50 × 22 complex points and 48 scans per free induction decay and two-dimensional ^15^N-TROSY experiments were collected with 512 × 64 complex points and 32 scans per free induction decay. The BEST-TROSY approach ^12^ was utilized in all experiments. Data were processed in NMRpipe^13^ and analyzed with POKY^14^ to generate 2D HNCO datasets from 3D HNCO data, processing of the carbonyl dimension was simply omitted and the first plane used. ^15^N-TROSY data were processed with the same parameters as the ^1^H and ^15^N dimensions of the 3D HNCO. Automated peak picking was conducted using UniDecNMR^2^. Peak centers were subsequently refined using the *voigt_fit* software in the DEEP Picker^1^ software suite. The resulting peak lists were then used directly without further curation for peak matching.

U-^2^H,^15^N,^13^C [MLV-^13^CH_3_] labeled His-tagged scFv of the fluorescein antibody 4-4-20 was expressed in *E. coli* BL21(DE3) cells during growth on minimal media containing 99% D_2_O supplemented with 2 g/L of deuterated glucose [^13^C-D7] and 1 g/L of ^15^NH_4_Cl (Cambridge Isotopes), isolated from inclusion bodies, and purified affinity chromatography on a HisTrap (Cytiva) nickel column. The protein was refolded by dropwise dilution. Details of the expression, purification and refolding will be presented elsewhere. The scFv:fluorescein complex was 0.2 mM in protein prepared in 10 mM sodium phosphate, 50mM NaCl at pH 6.0 with 5%(v/v) D_2_O, 0.02% (w/v) NaN_3_. Triple resonance experiments were carried out at 25°C (308K) on an 800 MHz(^1^H) four channel Bruker NEO NMR spectrometer equipped with a cryogenic probe. A summary of the experiments used is presented in Table 1. Chemical shift resonances were referenced with sodium 2,2-dimethyl-2-silapentane-5-sulfonate (DSS). Indirect dimension was acquired using NUS with Poisson gap distribution sampling and spectra were reconstructed using hmsIST^15^ and processed with NMRpipe in NMRBox^16^

**Table 1:** Summary of triple resonance spectra obtained for assignment of backbone resonances of the scFv of the fluorescein antibody 4-4-20.

|  |  |  |  |  |  |
| --- | --- | --- | --- | --- | --- |
|  | NS |  | <sup>1</sup>H (direct) | <sup>15</sup>N | <sup>13</sup>C |
| **HNCO** | 32 | pts | 2048 | 53 | 43 |
|  |  | SW (ppm) | 18.37 | 30.00 | 14.00 |
| **HN(CA)CO** | 96 | pts | 2048 | 53 | 43 |
|  |  | SW (ppm) | 18.37 | 30.00 | 14.00 |
| **HNCA** | 96 | pts | 2048 | 30 | 100 |
|  |  | SW (ppm) | 18.37 | 30.00 | 33.00 |
| **HN(CO)CA** | 48 | pts | 2048 | 30 | 100 |
|  |  | SW (ppm) | 18.37 | 30.00 | 33.00 |
| **HNCB** | 96 | pts | 2048 | 30 | 181 |
|  |  | SW (ppm) | 18.37 | 30.00 | 60.00 |
| **HN(CO)CB** | 64 | pts | 2048 | 30 | 181 |
|  |  | SW (ppm) | 18.37 | 30.00 | 60.00 |

### pHarmony parameters

For analysis of the spectra derived from the fragment-based ligand discovery screen of IL-1*β*, peak errors were set to 0.0075, 0.015, and 0.0015 ppm for ^13^C, ^15^N and ^1^H dimensions, respectively. The expected maximum CSP was set to 0.2 ppm and the expected fraction or reference cross-peaks undergoing a CSP to 0.1 with a variance scaling factor of 2. For analysis of the triple resonance spectra of the fluorescein ScFV, all of which were acquired on the same sample, the expected fraction of CSPs was set to 0.001. CSPs were calculated by scaling the chemical shifts in the carbon and nitrogen dimensions by a factor of 0.252 and 0.101 respectively before computing the Euclidian distance between cross-peaks.

## Results

### Harmonizing cross-peak sets anticipated to be closely identical

Perhaps the simplest and most efficient application of the pHarmony algorithm would be the rapid comparison of pairs of spectra containing common dimensional elements. This might include triple resonance spectra such as a ^15^N TROSY and HN(CO)CA where the spectra possess a set of axes reporting on identical resonances. Though relatively simple to manually digest, a rapid analysis that provides a statement of confidence for the matching of cross-peaks would significantly decrease the burden of manual analysis by highlighting those mappings of concern.

To evaluate this type of mapping, we employ a 30 kDa Fcsv construct of the fluorescein antibody 4-4-20 ^17^, which has been assigned using BARASA^18^. The assignments will be reported in detail elsewhere. This construct of 255 amino acids gives high quality triple resonance spectra as evidenced by the ^15^N-TROSY spectrum (Figure 2). Cross-peaks in the two-dimensional ^15^N-TROSY and three-dimensional triple resonance spectra (Table 1) were manually picked. The performance of pHarmony on this data set was evaluated using two criteria: accuracy (precision) and completeness (recall).

**Fig. 2.**
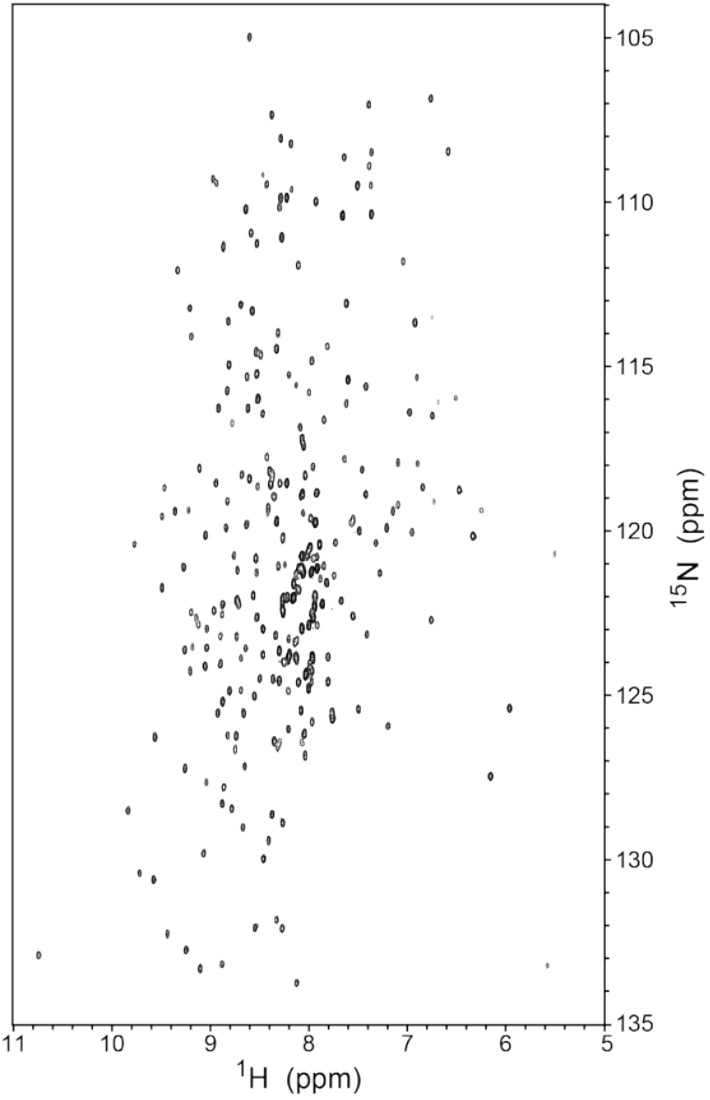
^15^N-TROSY spectrum of the scFv construct of the anti-fluorescein antibody 4-4-20 obtained at 800 MHz (^1^H) and 308 K.

Accuracy is defined as the ratio of “true” matches made by the algorithm to total matches made by the algorithm. Completeness is defined as the ratio of correct matches made by the algorithm to the total number of “true” matches. True matches were determined by manual annotation (where the triple resonance experiments provided unequivocal identification). Furthermore, since each match made is assigned a posterior probability, accuracy and completeness of matches can be evaluated as a function of match posterior probability, which serves as a confidence metric.

The performance of pHarmony when match triple resonance spectral pairs of the scFv construct of the anti-fluorescein antibody 4-4-20 is shown in Figure 3. A match was determined for each reference peak by selecting the highest probability matching target peak. These matches were then binned as a function of posterior probability. Employing a 95% posterior probability cutoff, the accuracy was 100% for the matches made in all cases. Completeness was in the range of 96-98%, meaning that the algorithm was missing very few of the possible matches at very high accuracy. The accuracy as a function of posterior probability did not appreciably follow the line of identity. However, this is likely due to high variance in the sampled accuracy over the low probability bins as the number of matches in posterior probability bins below 0.95 were all less than 6 per bin.

**Fig. 3.**
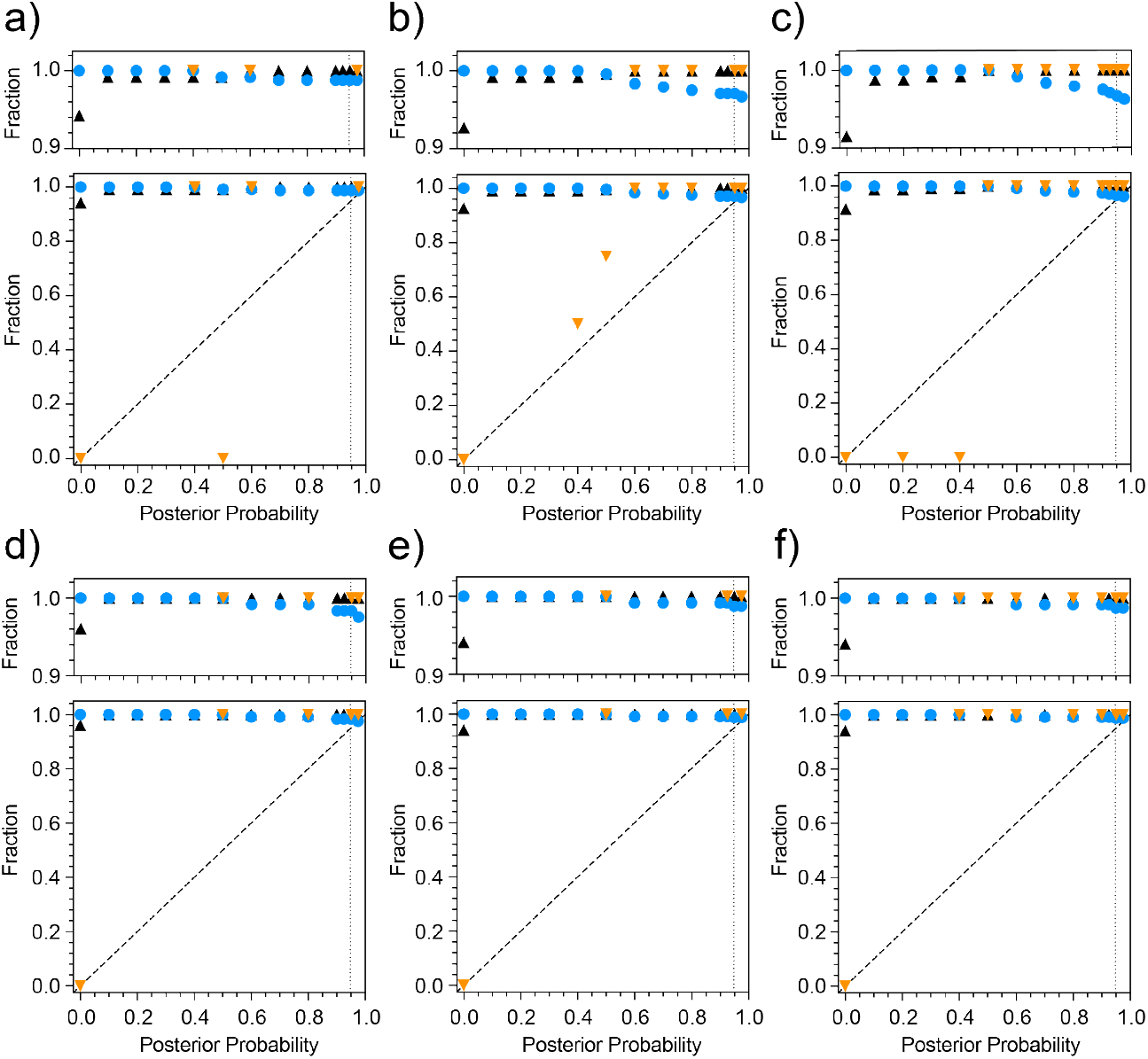
Performance of pHarmony in cross-peak matching between triple resonance spectra. Manually peak picked spectra of the 30 kDa anti-fluorescein scFv 4-4-20. Spectra were matched using only the chemical shift information from the ^15^N and ^1^H chemical shift coordinates. Shown for a given posterior probability are the accuracy of matches (orange triangles), cumulative completeness (blue circles) and cumulative accuracy (black triangles). Cumulative accuracy is defined as the accuracy of all matches at the indicated and all higher posterior probabilities. The accuracy data series omits points for empty posterior probability bins (i.e. no matches were made at that posterior probability range). Cross-peaks of the ^15^N-TROSY spectrum (reference) matched with ^1^H-^15^N dimensions of cross-peaks in the target HNCO (Panel a), HN(CO)CA (Panel b), HN(CO)CB (Panel c). The HNCO spectrum (reference) was matched with the target HN(CO)CA (Panel d) and HN(CO)CB (Panel e) spectra. The HN(CO)CA (reference) was matched with the target HN(CO)CB spectrum (Panel f). Spectra were matched using only the chemical shift information from the ^15^N and ^1^H chemical shift coordinates. The vertical dotted lines indicate a proposed confidence cutoff (0.95) for accepting matches.

### Harmonizing cross-peak sets anticipated to be significantly different

A more subtle and difficult task is the mapping of cross-peak sets that containing large numbers of chemical shifts differences. This situation can arise in numerous instances such as pH or other solvent changes, ligand titrations, temperature and pressure changes. To test pHarmony in this context, we utilized results from a fragment-based ligand discovery campaign of interleukin-1β (IL-1β) followed by multidimensional NMR spectra. This screen employed the reverse micelle encapsulation strategy to enhance detection of relatively weak binding fragments^10^. Details of the screen will be presented elsewhere. A rule-of-three (Ro3) fragment library customized for screening in the context of proteins encapsulated within reverse micelles (RM) was employed.

TROSY-HNCO and ^15^N-TROSY spectra were acquired on eleven samples of IL-1β encapsulated in 10MAG/LDAO reverse micelles. Each sample of IL-1β contained 5 different fragment compounds each present at an effective aqueous concentration of 40 mM. Two samples of IL-1β contained no fragments as a control. Each cross-peak in all spectra were manually annotated to serve as the ground truth. The carbon dimension of the HNCO was used to unambiguously track cross-peaks that moved in regions of degeneracy within the ^15^N-correlation (sub)spectrum. The algorithm was tested in four ways. The first three used manually picked peak lists and used to test 1) the ability to match cross-peaks between two HNCOs using all three dimension (3D-HNCO mode); 2) the ability to match cross-peaks between 2D HNCOs using the amide dimensions (2D HNCO mode); and 3) the ability to match cross-peaks between two HSQCs (2D-HSQC mode). The fourth test relied on automatically picked cross-peaks on HSQCs (2D-HSQC-Auto mode). In each mode, the algorithm was tested for its ability to match cross-peaks between all possible pairs of spectra (excluding matching a cross-peak list with itself). Additionally, since the algorithm treats reference and target cross-peaks differently, each pair of cross-peak lists was evaluated with each list as the target and the reference, leading to 110 runs for each mode.

The 3D-HNCO mode offers the opportunity to illustrate the distributions of normalized square distances of a more complex example than described in Figure 1. Figure 4 shows the three distribution classes, effectively noise and noise plus CSP matched distributions and the assumed unmatched function for comparison to two three-dimensional HNCO spectra of encapsulated IL-1β in the presence of two different mixtures of fragments.

**Fig. 4.**
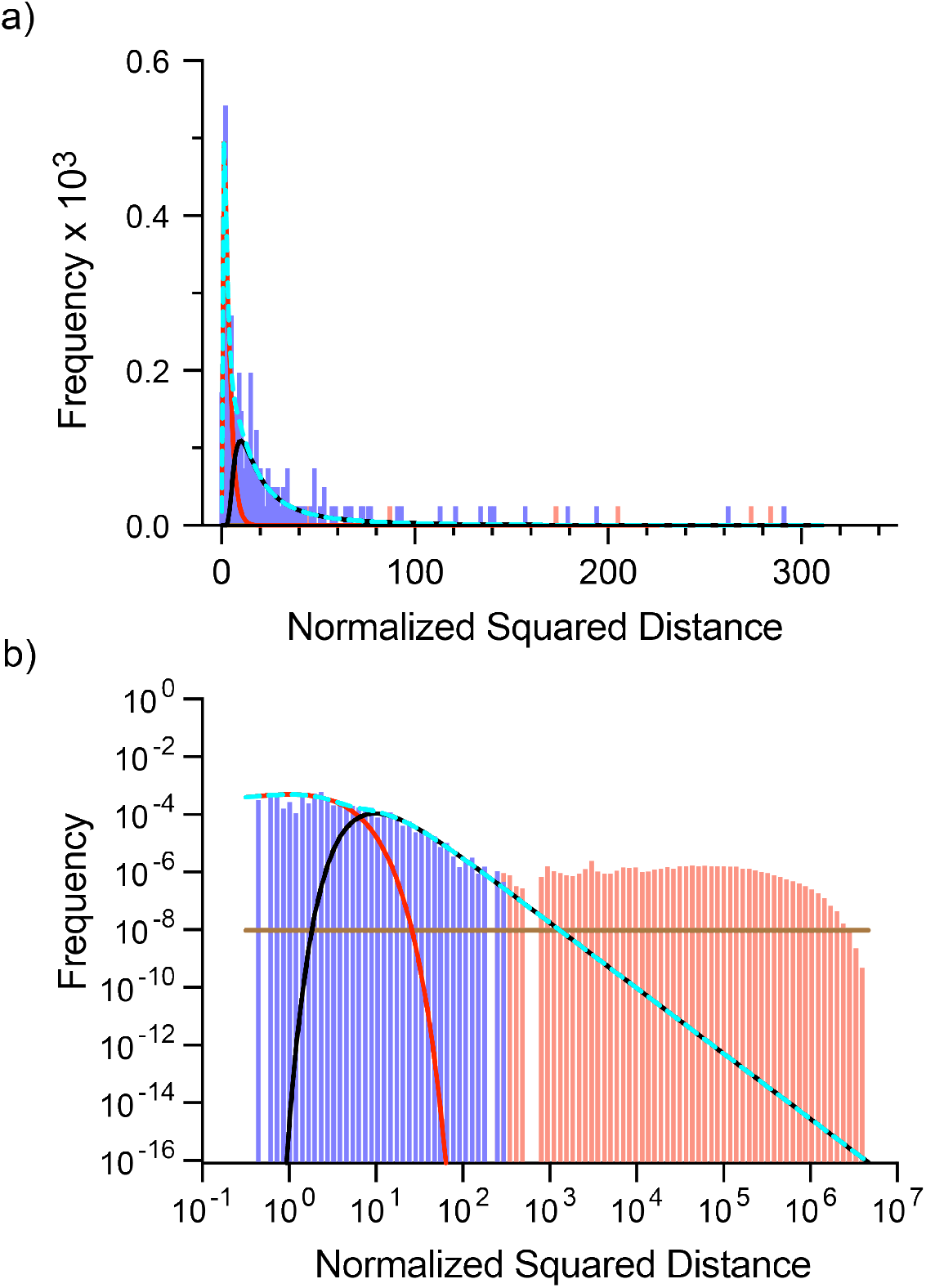
Example of distributions of normalized square distances of three-dimensional HNCO spectra. Pairwise normalized square distances between two IL1β 3D HNCO spectra were binned, with each distance classified as being either between peaks that match or do not match by the algorithm (at EM convergence). Blue-violet bars correspond to distances between matched peaks and red bars correspond to distances between peaks that were not matched. The solid red line corresponds to the component of the likelihood function for distances between matched peaks pair that did not undergo a CSP (Chi squared distribution). The solid black line corresponds to the component of the likelihood function for distances between matched peak pairs that undergo a CSP (Frechét distribution). The dashed cyan line corresponds to the matched probability distribution. The solid brown line corresponds to the pseudolikelihood function for distances between pairs of peaks that do not match. In this example, there are 198 reference peaks being compared to 207 target peaks giving 40,986 normalized squared distances. The top panel (a) is plotted in linear space showing the region containing all distances between matching pairs. The bottom panel (b) is plotted in log-log space showing all pairwise distances.

The performance of pHarmony for each of the testing modes is summarized in Figure 5. Among the testing modes involving manually picked cross-peaks, the 3D HNCO (Figure 5a) gave the best results both in terms of accuracy and completeness when considering the high posterior probability matches. The 2D HNCO and HSQC testing modes (Figure 5b & c, respectively) saw a slight decrease in the accuracy and completeness among the high posterior probability matches. However, if only accepting matches with an estimated posterior probability of 0.95 or greater, matches between cross-peaks of manually picked spectra have an accuracy of at least 98% and represent 86% of the “true” matchings. Performance using automatically picked cross-peaks on 2D HSQCs decreased both in terms of accuracy and completeness relative to any of the testing done with manually picked cross-peaks. Applying the with a posterior probability cutoff of 0.95 as before, completeness was at 71% of total matches with an accuracy of 94%.

**Fig. 5.**
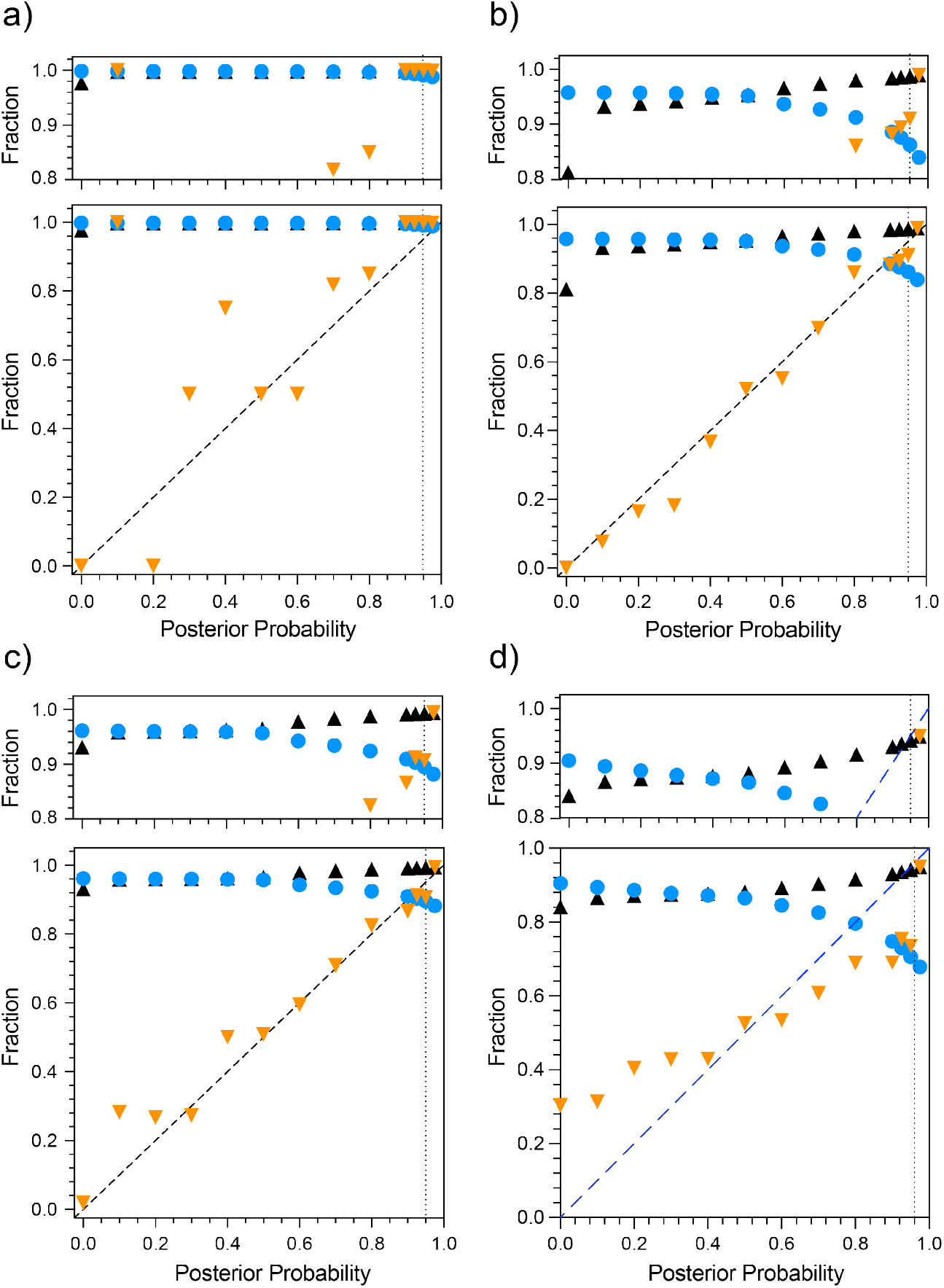
Performance of pHarmony across four different testing modes. Matches are binned by posterior probability. Matches of manually peak picked three-dimensional HNCO cross-peaks (Panel a), manually picked two-dimensional first ^13^C-increment of the HNCO cross-peaks (Panel b), manually picked two-dimensional ^15^N-HSQC cross-peaks (Panel c) and automatically picked two-dimensional ^15^N-HSQC cross-peaks (Panel d). Shown for a given posterior probability are the accuracy of matches (orange triangles), cumulative completeness (blue circles) and cumulative accuracy (black triangles). The vertical dotted lines indicate a proposed confidence cutoff for accepting matches.

The performance of pHarmony was compared to the peak matching algorithm Picasso which was designed to run on 2D spectra ^19^. Its performance was evaluated on the manually picked 2D HNCO, 2D HSQC and automatically picked 2D HSQC datasets using its implementation of the “RASmart” algorithm. The accuracy and completeness for each testing mode were (0.39, 0.46), (0.52, 0.53), and (0.40, 0.43), respectively.

There was a concern that the pHarmony algorithm could have difficulty in detecting larger CSPs. Though large CSPs tend to be less common in most contexts, cross-peaks that undergo large CSPs are often of the greatest interest. By the nature of the probability model, the likelihood of two cross-peaks matching decreases as the distance between them increases. The algorithm eventually begins to favor matches to other, closer cross-peaks or possibly matching no peak at all. Figure 6 shows the distributions of “detected” true matches (correct matches with a posterior probability of 0.95 or greater) and “missed” true matches (i.e., true matches that were either made by the algorithm with a posterior probability below the cutoff or entirely undetected by the algorithm) binned by CSP for each of the testing modes. Peak matching manually picked cross-peaks in the 3D HNCO had the best results in terms of detecting long range CSPs, with the 2D HNCO and 2D HSQC being less capable of their detection. Peak matching automatically picked HSQCs had the lowest performance with the algorithm being more likely to miss true matches than detect them beyond CSPs of 0.025 ppm.

**Fig. 6.**
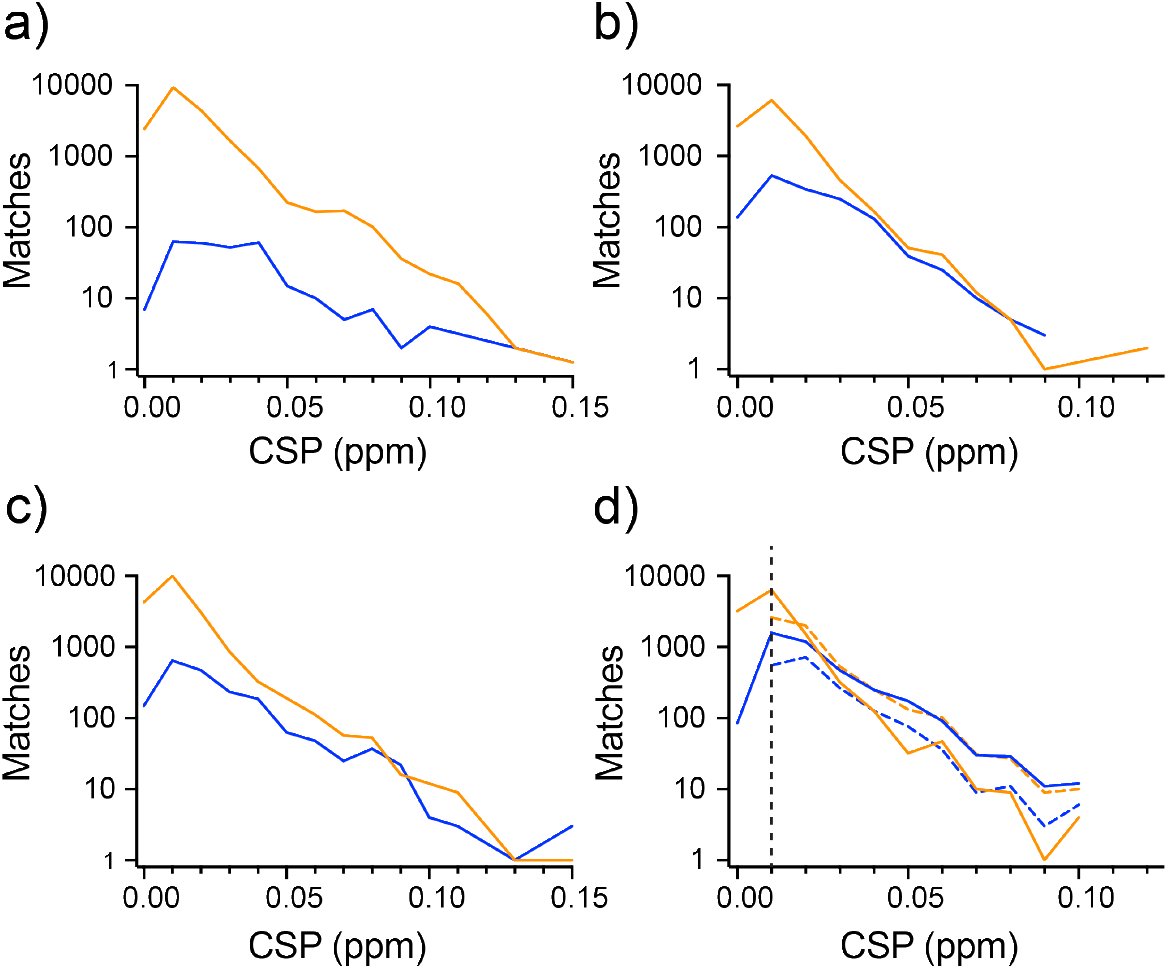
Success of pHarmony cross-peak matching between spectra in the presence of chemical shift perturbations. Semi-log plots of numbers of matches made of manually picked cross-peaks between the 3D-HNCO spectra (Panel a), those of the ^1^H-^15^N plane of the first carbon increment of the 3D-HNCO spectra (Panel b), and those of the ^15^N-TROSY spectra (Panel c). Panel d compares matching of automatically picked cross-peaks of the ^15^N-TROSY spectra. Orange and blue solid lines represent matches made with greater than or less than 0.95 posterior probability, respectively. Dashed lines represent the numbers of found and missed CSPs when the integrated marginal posterior probability over target peak matches that each correspond to a significant CSP was greater than 0.95. The vertical dashed line corresponds to the cutoff used to define a significant CSP. See text for details.

In the context of a ligand screening campaign, large CSPs are often the most valuable to identify as they are potentially indicative of compounds of higher affinity. It is in this context that a probabilistic approach has considerable value. It is not necessary that a hit detection algorithm confidently identify a single, definite match for a reference peak in a target spectrum to determine whether a CSP has occurred. Rather, if the algorithm could confidently identify that a reference peak matches any from a subset of target cross-peaks, all of which correspond to a “significant” CSP, then the user could confidently assert that a peak was significantly perturbed even if there is some uncertainty as to how it was perturbed. Answering this question for a given reference spectrum peak requires its marginal matching probability distribution over the possible target cross-peaks. As the algorithm samples the *joint* matching probability distribution, estimation of the marginal distribution involves integrating over the sampled matching matrices to determine the frequency with which the reference peak in question matched each target peak. If a large percentage (>95%) of the marginal probability mass corresponded to target peak matches above a CSP significance cutoff (> 0.01 ppm). Then the user could safely conclude that the peak was perturbed above the CSP significance cutoff without exactly knowing which target peak it matched. When employing this strategy of CSP detection (Figure 6d) there was a substantial improvement in the sensitivity of the algorithm for large CSP detection. The accuracy of the detected CSPs using this strategy was 0.94 with the completeness being 0.76.

## Discussion

The pHarmony algorithm was able to obtain a high degree of accuracy and completeness on matches between both 2D and 3D spectra when employing manually picked cross-peaks. Performance in terms of both accuracy and completeness was better for the 3D spectra likely due to the extra resolution afforded by the third dimension. Furthermore, the posterior probability of the matches was very close to the probability that the resulting match was correct as shown by the tendency of the accuracy to track the line of identity (see Figure 5). This demonstrates that the computed posterior probability is a reliable indicator of confidence in the veracity of a proposed match. Though accuracy as a function of posterior probability for the 3D HNCO test case has quite a bit of scatter, it should be noted that the number of matches in each bin below posterior probability 0.5 was less than 10. These small sample sizes likely contributed to high variance in the estimated accuracy at low posterior probabilities. It also supports the validity of the model used to compute matching likelihoods, as it demonstrates that the calculated posterior probability closely approximates the quantity that it is trying to model which is the probability a proposed match between a reference peak and target peak is correct.

The completeness of the matches at high accuracy experienced a substantial drop when utilizing automatically picked cross-peaks compared to its manually picked counter parts. We ascribe this occurrence to circumstances where the automatic peak picking routine picks a peak in a crowded region of the reference spectrum, but does not pick it in the target spectrum (or vice-versa). If the same peak is not picked in the target spectrum, then the presence of the extra reference peak reduces confidence in otherwise confident nearby matches. In addition, the accuracy as a function of posterior matching probability did not track the line of identity as well. However, it is important to note that the cumulative accuracy of the algorithm at the posterior probability cutoff of 0.95 was 94%. Thus, the posterior probability still provides a reliable prediction of accuracy at high confidence cutoffs.

In the case of the manually picked cross-peaks, the chance of missing a true match was only weakly related with the size of the corresponding chemical shift perturbation. Though the probability of finding a reference peak’s match is best when it lies in its immediate vicinity (< 0.02 ppm). The chance of finding the matching peak is relatively constant as a function of increasing chemical shift from approx. 0.2 ppm to the largest CSPs in the data set (approx. 0.1 ppm). This is indicated by the similar slope (in log space) of the distributions of found and missed matches in Figure 6. This was true in all test conditions involving manually picked cross-peaks. In manually picked 2D spectra, the algorithm was equally likely to find a CSP greater than 0.03 ppm as it was to miss it. This suggests that the reasons for the algorithm missing cross-peaks lies more in ambiguity due to spectral crowding than distance. In the automatically picked HSQC cases, there is relationship between the CSP of a valid match and the ability of the algorithm to detect it. At higher CSPs (> 0.025 ppm) the algorithm becomes more likely to miss a true match than detect it. This is possibly due to additional ambiguity introduced from cross-peaks being erroneously picked or missed by automatic peak picking.

If the goal is to use automated peak picking for the purposes of hit detection, computation of the overall CSP probability for each reference peak from their respective marginal matching probability distributions resulted in substantial improvements in sensitivity. The completeness of the found CSPs increased from 0.58 (when requiring a 0.95 posterior probability match to a single target peak greater than 0.010 ppm away) to 0.76 (when requiring a cumulative posterior probability of at least 0.95 summed over all target cross-peaks more than 0.010 ppm away). The accuracy did not change appreciably (0.95 to 0.96). Importantly this improvement in completeness was achieved through significantly better detection of long range CSPs, which are often of greatest interest in a screening context. It is even possible to get an estimate of the CSP magnitude and uncertainty when considering multiple high probability matches by taking a marginal probability weighted average and standard deviation of the CSPs over all possible target cross-peaks.

## Conclusions

A novel algorithm (pHarmony) for tracking peak movement due to a perturbation between two spectra has been described. pHarmony fills a gap left by existing tools: prior efforts emphasize automated peak picking (e.g., DEEP picker^1^, UniDecNMR^2^, CYPICK^3^) or 2D-only CSP matching (e.g., PICASSO^19^), typically without formal confidence estimates or support for general multidimensional peak lists. The use of modern probabilistic machinery (mixture modeling, EM, SMC, beam search) is well aligned with current trends in quantitative NMR analysis. Cross-peak matching subject to random noise with and without chemical shift changes due to an external perturbation are well handled. While ligand binding was the perturbation tested here, the algorithm could be used for any kind of perturbation inducting peak movement (e.g., temperature, pH, salt, etc.). The algorithm natively handles situations where cross-peaks may disappear either due to experimental phenomena (e.g., intermediate exchange) or data analysis errors (e.g., erroneous peak picking) and can work on peak lists of arbitrary dimensionalities. Importantly, the algorithm provides a robust confidence estimate of proposed matches that closely approximates true accuracy of the algorithm. In addition, the python package contains a peak matching function that can be imported into user-written python scripts for seamless integration into automated workflows providing direct access to the estimated marginal matching distributions of the reference and target cross-peaks.

## Software Availability

The python package containing pHarmony is accessible through NMRbox and can be downloaded from https://github.com/anthony-bishop-tamu/pHarmony. pHarmony was written employing the numpy^20^, scipy^7^, matplotlib^21^, pandas^22^, pytorch^23^, scikit-learn^6^, and tqdm^24^ python packages as dependencies.

## Acknowledgments

This work was supported by the NIH (R01 GM145751 and R35 GM158127), the Welch Foundation (A-2258-20250403) and Texas A&M University.

## References

1. Li, D.W., Hansen, A.L., Yuan, C., Bruschweiler-Li, L. & Bruschweiler, R. DEEP picker is a deep neural network for accurate deconvolution of complex two-dimensional NMR spectra. Nature Communications 12, 5229 (2021).

2. Buchanan, C. et al. UnidecNMR: automatic peak detection for NMR spectra in 1-4 dimensions. Nature Communications 16, 449 (2025).

3. Wurz, J.M. & Guntert, P. Peak picking multidimensional NMR spectra with the contour geometry based algorithm CYPICK. Journal of Biomolecular NMR 67, 63–76 (2017).

4. Jun, S.H., Wong, S.W.K., Zidek, J.V. & Bouchard-Côté, A. Sequential Graph Matching with Sequential Monte Carlo. Journal of Machine Learning Research 54, 1075–10184 (2017).

5. Floyd, R.W. Algorithm 97: Shortest path. Communications of the ACM 5, 345–345 (1962).

6. Pedregosa, F. et al. Scikit-learn: Machine Learning in Python. Journal of Machine Learning Research 12, 2825–2830 (2011).

7. Virtanen, P. et al. SciPy 1.0: fundamental algorithms for scientific computing in Python. Nature Methods 17, 261–272 (2020).

8. Ow, P.S. & Morton, T.E. Filtered beam search in scheduling. International Journal of Production Research 26, 35–62 (1988).

9. Kitagawa, G. Monte Carlo Filter and Smoother for Non-Gaussian Nonlinear State Space Models. Journal of Computational and Graphical Statistics 5, 1–25 (1996).

10. Fuglestad, B., Kerstetter, N.E., Bedard, S. & Wand, A.J. Extending the Detection Limit in Fragment Screening of Proteins Using Reverse Micelle Encapsulation. ACS Chemical Biology 14, 2224–2232 (2019).

11. Dodevski, I. et al. Optimized reverse micelle surfactant system for high-resolution NMR spectroscopy of encapsulated proteins and nucleic acids dissolved in low viscosity fluids. Journal of the American Chemical Society 136, 3465–74 (2014).

12. Favier, A. & Brutscher, B. Recovering lost magnetization: polarization enhancement in biomolecular NMR. Journal of Biomolecular NMR 49, 9–15 (2011).

13. Delaglio, F. et al. NMRPipe: a multidimensional spectral processing system based on UNIX pipes. Journal of Biomolecular NMR 6, 277–93 (1995).

14. Lee, W., Rahimi, M., Lee, Y. & Chiu, A. POKY: a software suite for multidimensional NMR and 3D structure calculation of biomolecules. Bioinformatics 37, 3041–3042 (2021).

15. Hyberts, S.G., Milbradt, A.G., Wagner, A.B., Arthanari, H. & Wagner, G. Application of iterative soft thresholding for fast reconstruction of NMR data non-uniformly sampled with multidimensional Poisson Gap scheduling. Journal of Biomolecular NMR 52, 315–27 (2012).

16. Maciejewski, M.W. et al. NMRbox: A Resource for Biomolecular NMR Computation. Biophys J 112, 1529–1534 (2017).

17. Kranz, D.M. & Voss, E.W., Jr. Partial elucidation of an anti-hapten repertoire in BALB/c mice: comparative characterization of several monoclonal anti-fluorescyl antibodies. Molecular Immunology 18, 889–98 (1981).

18. Bishop, A.C., Torres-Montalvo, G., Kotaru, S., Mimun, K. & Wand, A.J. Robust automated backbone triple resonance NMR assignments of proteins using Bayesian-based simulated annealing. Nature Communications 14, 1556 (2023).

19. Laveglia, V. et al. Automated Determination of Nuclear Magnetic Resonance Chemical Shift Perturbations in Ligand Screening Experiments: The PICASSO Web Server. Journal of Chemical Information and Modeling 61, 5726–5733 (2021).

20. Harris, C.R. et al. Array programming with NumPy. Nature 585, 357–362 (2020).

21. Barrett, P., Hunter, J., Miller, J.T., Hsu, J.C. & Greenfield, P. matplotlib - A portable python plotting package. Astronomical Data Analysis Software and Systems XIV, Proceedings 347, 91–95 (2005).

22. McKinney, W. Data structures for statistical computing. Proceedings of the 9th Python in Science Conference 445, 56–61 (2010).

23. Paszke, A. et al. PyTorch: An Imperative Style, High-Performance Deep Learning Library. Advances in Neural Information Processing Systems 32 (NeurIPS 2019) 32 (2019).

24. Costa-Luis, d. et al. tqdm: A fast, Extensible Progress Bar for Python and CLI. (Zenodo, 2026).

